# Causal inference explains the stimulus-level relationship between the McGurk Effect and auditory speech perception

**DOI:** 10.1101/2020.05.08.085209

**Authors:** John F. Magnotti, Kristen B. Dzeda, Kira Wegner-Clemens, Michael S. Beauchamp

## Abstract

The McGurk effect is widely used as a measure of multisensory integration during speech perception. Two observations have raised questions about the relationship between the effect and everyday speech perception. First, there is high variability in the strength of the McGurk effect across different stimuli and observers. Second, there is low correlation across observers between perception of the McGurk effect and measures of everyday speech perception, such as the ability to understand noisy audiovisual speech. Using the framework of the causal inference of multisensory speech (CIMS) model, we explored the relationship between the McGurk effect, syllable perception, and sentence perception in seven experiments with a total of 296 different participants. Perceptual reports revealed a relationship between the efficacy of different McGurk stimuli created from the same talker and perception of the auditory component of the McGurk stimuli presented in isolation, either with or without added noise. The CIMS model explained this high stimulus-level correlation using the principles of noisy sensory encoding followed by optimal cue combination within a representational space that was identical for McGurk and everyday speech. In other experiments, CIMS successfully modeled low observer-level correlation between McGurk and everyday speech. Variability in noisy speech perception was modeled using individual differences in noisy sensory encoding, while variability in McGurk perception involved additional differences in causal inference. Participants with all combinations of high and low sensory encoding noise and high and low causal inference disparity thresholds were identified. Perception of the McGurk effect and everyday speech can be explained by a common theoretical framework that includes causal inference.

## Introduction

Viewing the talker’s face influences the perception of auditory speech, as exemplified by McGurk and MacDonald’s discovery that pairing incongruent auditory and visual syllables can evoke the percept of a completely different syllable (McGurk and MacDonald, 1976). In the decades since its discovery, the McGurk effect has grown into one of the most popular experimental tools for assessing multisensory integration, with thousands of citations across both behavioral and neural sciences (Beauchamp, 2018).

Recently, doubts have arisen about the utility of the McGurk effect as a tool for understanding everyday forms of speech perception, including the suggestion that it should be “retired” (Rosenblum, 2019). In a large study, Basu Mallick and colleagues examined perception of 12 different McGurk stimuli by 165 participants tested in the laboratory (Basu Mallick et al., 2015). There was high variability, both across stimuli (rates ranging from 17% to 58%) and across participants (rates from 0% to 100%). This high variability has been used as an argument that the McGurk effect is unreliable and hence poorly suited for experimental study (Alsius et al., 2018).

A second critique of the McGurk effect is the difficulty of relating it to everyday speech perception. Viewing the talker’s face benefits understanding of noisy auditory speech (Sumby and Pollack, 1954; Peelle and Sommers, 2015). Van Engen and colleagues (Van Engen et al., 2017) found high variability in visual enhancement of noisy speech and in rates of the McGurk effect but there was low correlation between the two measures for any of the twelve types of noisy speech examined, with a maximum of 4% variance explained. Similarly, Brown and colleagues (Brown et al., 2018) found only a weak correlation between lipreading accuracy and McGurk susceptibility (3% of variance explained; 8% variance if lipreading responses were grouped by place-of-articulation).

We reasoned that modeling the processes underlying audiovisual speech perception might shed light on these observations and the relationship between the McGurk effect and everyday speech in general. Two processes commonly assumed to underlie all types of multisensory perception are *noisy sensory encoding* and *optimal cue combination*. Noisy sensory encoding assumes that observers to do not have veridical access to the physical properties of a stimulus, but only to a perceptual representation that is corrupted by sensory noise and that can vary on repeated presentations of identical stimuli (Deneve et al., 2001). Optimal cue combination assumes that when cues from different modalities are combined, they are weighted by the reliability of each modality, a process often referred to as Bayesian integration (Ernst and Banks, 2002; Alais and Burr, 2004; Magnotti et al., 2013; Aller and Noppeney, 2019) although there are alternative algorithms such as probability summation (Arnold et al., 2019). Optimal cue combination also requires *causal inference*, judging whether the cues in the different modalities arise from the same physical cause. Causal inference is necessary because it is only beneficial to integrate cues generated by the same source; integrating cues from different sources leads to misestimation (Kording et al., 2007; Shams and Beierholm, 2010; French and DeAngelis, 2020). The necessity of causal inference in audiovisual speech perception has been widely recognized (Massaro, 1998; Ma et al., 2009; Vroomen, 2010; Noppeney and Lee, 2018) and individual differences in causal inference judgments have been used to characterize individual- and group-level differences in audiovisual speech perception (Magnotti et al., 2013; Baum et al., 2015; Gurler et al., 2015; Magnotti and Beauchamp, 2015; Stropahl et al., 2017).

The causal inference of multisensory speech (CIMS) model incorporates these processes into a principled framework that predicts perception of arbitrary combinations of auditory and visual speech (Magnotti and Beauchamp, 2017). The CIMS model has been used to explain a number of puzzling audiovisual speech phenomena, such as the increase in the McGurk effect observed with co-articulation (Magnotti et al., 2018b); the decrease in the McGurk effect observed with slower playback rates (Magnotti et al., 2018a); and why the McGurk effects is produced by some incongruent syllables but not others (Magnotti and Beauchamp, 2017). If the CIMS model (or any other model) can account for perception of McGurk and everyday speech by explaining high variability across stimulus and participant and low correlation between McGurk and other measures of speech perception, it suggests that McGurk and everyday speech may be processed using common perceptual mechanisms, with the implication that the McGurk effect is a useful tool for interrogating everyday speech processing. On the other hand, if CIMS model predictions derived from the McGurk effect do not apply to everyday speech, it suggests that that the McGurk effect has limited utility for understanding everyday speech processing (Alsius et al., 2018; Rosenblum, 2019).

## Methods

### Human Subject Statement

All experiments were approved by the Committee for the Protection of Human Subjects of Baylor College of Medicine.

### Overview of Experimental Procedures

Participants viewed brief recordings of audiovisual speech and reported their percepts. Experiments 1 - 6 examined perception of syllables (both McGurk and congruent) using the online data collection service Amazon Mechanical Turk (Buhrmester et al., 2018). In a previous study, we found that online testing gives similar results as in-person testing (Magnotti et al., 2018b). Experiment 7 examined the McGurk effect and perception of noisy sentences, with data collected in-person at Baylor College of Medicine. A total of 262 different participants were tested online and 34 different participants were tested in person for a total of 296 different participants.

All data was analyzed using R (Computing, 2017). Variability was modeled using linear mixed effects models (LMEs) as implemented in the lme4 packager (Bates et al., 2015). LMEs provide a consistent approach for understanding the effect of both categorical and numeric independent variables (fixed effects) while taking into account other sources of variation (random effects such as participant effects or stimulus effects). To test the significance of the fixed effects, we report *t*-tests with Satterthwaite-approximated degrees of freedom, as implemented in the lmerTest package (Kuznetsova et al., 2017).

### Online Testing Procedures for Experiments 1 - 6

Participants enrolled as workers in the Mechanical Turk service and requested to participate in our experiment, in return for compensation of approximately $10 per hour. After accepting the assignment, they were directed to an informed consent statement, followed by completion of a demographic questionnaire. Before starting the experiment, participants were shown a demonstration video. Participants were instructed to resize their browser window and their computer audio as needed to make the demonstration video easily visible and audible. The demonstration video could be repeated as often as necessary by the participant. Then, the participants proceeded to the main experiment, which consisted of multiple trials. Within each trial, participants viewed a single stimulus and reported their percept using a forced-choice response, selecting among three possibilities, “ba” (the auditory component of the stimulus), “ga” (the visual component of the stimulus) or “da/tha” (the McGurk or fusion percept). In a previous study, we demonstrated similar results between this set of forced-choice responses and open-choice responding (Basu Mallick et al., 2015). Forced-choice has the advantage of reducing the time to analyze data and of reducing the experimenter degrees of freedom available when quantifying open-choice responses. No feedback was given to reduce demand characteristics.

### Experiment 1 Stimulus Creation

The experimental stimulus set in Experiment 1 consisted of twenty different McGurk syllables (auditory /ba/ paired with visual /ga/, A/ba/V/ga/) where the /ba/ was different in each syllable but the /ga/ was identical. A female native speaker of American English was recorded voicing the syllable /ba/ twenty times. The auditory component of each stimulus was imported into MATLAB and the volume of each clip was normalized by dividing the sound amplitude by the square root of the squared mean. The resulting waveform was scaled to prevent clipping, with each auditory track scaled to the same power.

The same talker was recorded saying a single /ga/ using a Panasonic AG-HVX200AP video camera. The camera view showed the talker’s head and shoulders against a white background. The video obtained was imported into Adobe Premiere Pro CC 2015. Then, each of the twenty auditory /ba/ recordings was dubbed onto the visual portion of the /ga/ recording so that the auditory and visual components were synchronized. Each was exported to QuickTime format and Handbrake software was used to crop and convert the videos to 640 by 480 resolution in the MP4 format.

The control stimulus set in Experiment 1 consisted of audiovisual recordings of a different female native speaker of American English speaking three syllables for which the auditory and visual components were congruent, (A/ba/V/ba/, A/ga/V/ga/, A/da/V/da/).

To avoid participant fatigue or adaptation, each participant was presented with five different McGurk stimuli (randomly selected from the entire battery of twenty) and the three congruent stimuli. Each McGurk stimulus was presented nine times and each congruent stimulus was presented three times, all randomly interleaved.

A total of 160 participants were recruited (57 female, 93 male, 10 did not specify). The 20 stimuli were tested in 4 batches of 5 stimuli each. The first 40 participants viewed the first five stimuli; the second 40 participants viewed the next five stimuli; and so on. Since each participant viewed one-quarter of the stimuli (5 / 20), the final N was 40 participants for each of the twenty McGurk stimuli.

### Experiment 2

From the twenty different McGurk stimuli presented in Experiment 1, we selected two stimuli at opposite ends of the distribution for further investigation, labeling them “S1” and “S2”. A total of 40 participants were recruited for experiment 2 (10 female, 25 male, 5 did not specify). Each participant was presented with 9 repetitions each of S1 and S2 and three repetitions of the congruent stimuli, randomly interleaved, and participants made forced choice responses.

### Experiment 3

Praat (Boersma and Weenink, 2020) was used to create a morphed stimulus intermediate between S1 and S2, labeled “S1.5”. As the visual /ga/ component of S1 and S2 was identical, only the auditory component of S1.5 differed from S1 and S2. The experimental stimulus set in Experiment 3 consisted of the auditory-only /ba/ component of S1, S1.5 and S2. The control stimulus set consisted of the auditory-only component of the control stimuli in Experiment 1, auditory /ba/, /da/ and /ga/. The visual component for all stimuli consisted of white text on a black square instructing participant to “Listen to the audio.” 40 participants (14 female, 23 male, 3 did not specify) were presented with nine repetitions of each of the experimental stimuli and three repetitions of each of the control stimuli, all randomly interleaved.

### Experiment 4

The experimental stimuli were the audiovisual A/ba/V/ga/ syllables S1, S1.5 and S2. The control stimuli were the congruent syllables A/ba/V/ba/, A/ga/V/ga/ and A/da/V/da/. 40 participants (20 female, 19 male, 1 did not specify) were presented with nine repetitions of each McGurk stimulus and three repetitions of each congruent syllable, all randomly interleaved.

### Experiment 5

Auditory noise was added to S1 and S2 by combining each stimulus with uniform white noise with signal to noise ratios (SNRs) of -30, -24, -18, -12, 0 dB and no noise. Final Cut Pro was used to create the noisy stimuli by combining the noise clip and the syllable clip and adjusting the decibel levels of each to the desired SNR. After noise was added, the volume of each clips was RMS normalized in MATLAB and the stimuli were imported into Final Cut Pro to add a blank visual screen for the visual component, followed by resizing in Handbrake to a 640:480 aspect ratio. 44 participants (21 female, 22 male, 1 did not specify). Each of the two stimuli (the /ba/ extracted from S1 and S2) was presented six times at each of the six noise levels (72 total presentations). Control stimuli consisted of two examples of auditory /da/ and two examples of auditory /ga/ recorded by the same talker. The same six noise levels were added to each control stimulus and each was presented six times, ensuring an equal total number of experimental and control stimulus presentations (72 for each).

### Experiment 6

The 44 participants from Experiment 5 were invited to return. 38 participants returned (18 female, 19 male, 1 did not specify) and rated the McGurk stimuli S1, S1.5 and S2 to allow for intra-participant comparison of noisy syllable and McGurk perception. Experimental stimuli consisted of 10 presentations each of the McGurk stimuli S1, S1.5 and S2. Control stimuli consisted of three presentations each of the congruent audiovisual syllables A/ba/V/ba/, A/ga/V/ga/ and A/da/V/da/.

### Experiment 7 Overview

34 native English speakers (18 female, 16 male) were tested in-person at the Core for Advanced MRI at Baylor College of Medicine. All tasks were presented using Matlab (Mathworks, Inc., Natick, MA, USA) with the Psychophysics Toolbox extensions (Brainard, 1997; Pelli, 1997). Visual speech was presented on a high-resolution screen (Display++ LCD Monitor, 32-in., 1,920 × 1,080, 120 Hz, Cambridge Research Systems, Rochester, UK). Auditory speech was presented through speakers on either side of the screen at a constant sound pressure level of 60 dB, a value chosen to approximate the level of human speech.

### Experiment 7: McGurk and congruent syllables

First, participants viewed stimuli consisting of 2-second recordings of audiovisual syllables with no added noise. The syllables were recorded by five different talkers. Each participant was presented with a total of 270 trials: 20 repetitions × four talkers × three syllables (two congruent: A/ba/V/ba/, A/da/V/da/; one McGurk A/ba/V/ga/) and 10 repetitions × one talker × three audiovisual syllables (all congruent: A/ba/V/ba/, A/da/V/da/,A/ga/V/ga/), all randomly interleaved. Participants reported the identity of each syllable (“ba”, “da”, or “ga”) with a button press.

### Experiment 7: Noisy speech

Second, participants were presented with 3-second duration sentences recorded from a single male talker combined with auditory pink noise at a signal-to-noise ratio (SNR) of -16 dB, as used in a previous study (Van Engen et al., 2017). The sentences were presented either alone (noisy auditory-only, A) or paired with a video recording (noisy auditory + visual, AV). After the sentence ended, participants repeated the sentence aloud. Responses were scored for number of correct keywords (e.g., “The **hot sun warmed** the **ground**,” keywords in bold). Each participant was presented with 80 sentence trials, consisting of randomly interleaved presentations of 40 auditory-only and 40 audiovisual sentences. To prevent perceptual learning, individual sentences were never repeated.

## Results

### Section 1: Stimulus Variability

If McGurk and everyday speech are processed using common perceptual mechanisms, then variability across stimuli should create predictable changes in both McGurk and everyday perception. For instance, even for a single talker, there is substantial variability in repeated productions of the same speech token (Holmberg et al., 1994; Whalen et al., 2018). In CIMS, this variability is modeled with a representational space that collapses all auditory and visual speech features onto a one-dimensional line with /ba/, /da/, and /ga/ at neighboring locations. One production of the syllable /ba/ (Stimulus 1) might lie near the protoypical /ba/, while a second production (Stimulus 2) might lie further from the prototype (Fig. 1A).

**Figure 1.**
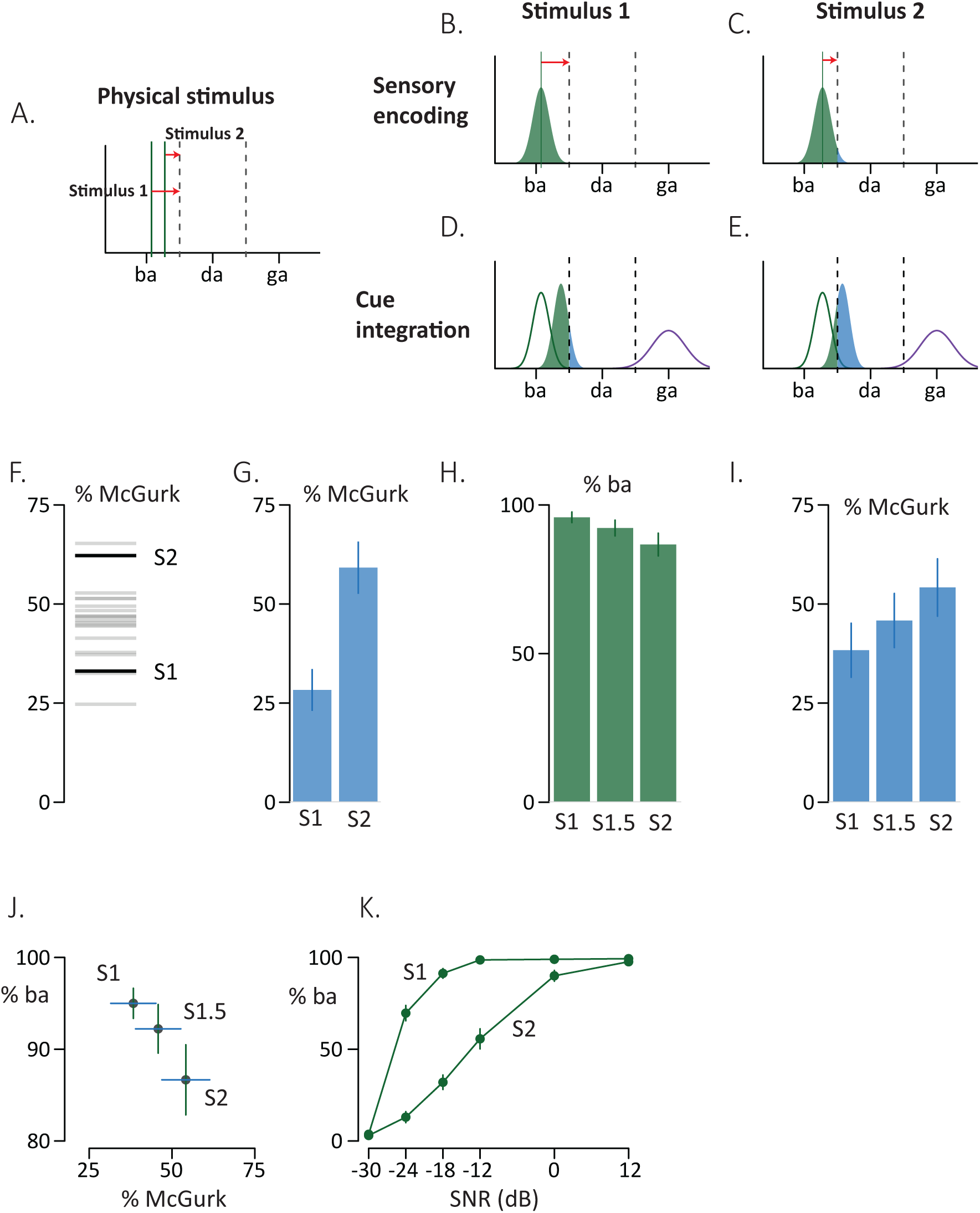
The CIMS model applied to stimulus variability. (A) In the causal inference of multisensory speech perception (CIMS) model, the physical properties of auditory and visual speech can be collapsed onto a one-dimensional representational space with different regions of the space representing different tokens (/ba/, /da/ and /ga/ shown). The physical properties of a token determine its location in representational space, as shown for two example /ba/ tokens. The left token (Stimulus 1) is closer to the center of the /ba/ region of representational space and hence is a more prototypical /ba/. The right token (Stimulus 2) is further from the center of the /ba/ region of representational space, and hence is a less prototypical /ba/. Dashed lines indicate boundaries between different regions of representational space. (B) In the sensory encoding stage of the CIMS model, the physical properties of the stimulus are encoded with sensory noise, resulting in a distribution of encoded values for any given token. The mean of the distribution is determined by the physical properties of the stimulus and the variance of the distribution is determined by the sensory noise for that modality for that observer. For Stimulus 1, the stimulus is far from the perceptual boundary (distance indicated by red arrow), with the result that even after noisy encoding, most perceived locations are in the /ba/ region of representational space, resulting in exclusively /ba/ percepts (green shaded region). (C) For Stimulus 2, the stimulus is close to the perceptual boundary (distance indicated by red arrow) with the result that after noisy encoding, some perceived locations are in the /da/ region of representational space (blue shaded region). (D) In the cue integration stage of the CIMS model, auditory cues (green lines) and visual cues (purple lines) are integrated using optimal cue combination, resulting in an average representation weighted by the reliability of each modality. For Stimulus 1, most perceived locations for the integrated representation are in the /ba/ region of representational space, resulting in mainly /ba/ percepts (green shaded region). (E) For Stimulus 2, the location of the auditory component nearer the perceptual boundary means that many perceived locations for the integrated representation are in the /da/ region of representational space, the McGurk fusion percept (blue shaded region). (F) In experiment 1, twenty different /ba/ tokens recorded by the same talker were paired with a single /ga/ from the same talker to create twenty different AbaVga McGurk stimuli. Each bar represents the % of McGurk fusion for a single stimulus. Two stimuli were selected for further analysis, dark bars labelled “S1” and “S2”, analogous to modeled Stimulus 1 and Stimulus 2. (G) In experiment 2, S1 and S2 were presented to a different set of participants. There was no significant difference between experiment 1 and the replication sample in experiment 2 (*p* = 0.66). (H) In experiment 3, a morphed stimulus was created by combining S1 and S2 (labeled “S1.5”) and the auditory-only /ba/ component of S1, S1.5 and S2 was presented. Mean and SEM of % “ba” percepts for each stimulus are shown. (I) In experiment 4, the McGurk stimuli S1, S1.5 and S2 were presented, mean and SEM of % McGurk percepts for each stimulus are shown. (J) The rates of McGurk perception for S1, S1.5 and S2 were plotted against the perceptual accuracy for the auditory component of each stimulus, (I) *vs.* (H). (K) In experiment 5, different levels of auditory noise were added to the auditory components of S1 and S2 and the rate of “ba” responses measured.

An important feature of the CIMS model is sensory encoding noise. While the physical properties of a given speech token place it at one location in representational space, sensory encoding is noisy, with the result that repeated presentations of the identical token produce a distribution of perceived locations whose mean is at the true location and whose variance is proportional to the amount of sensory noise. Over repeated presentations of Stimulus 1, its location far from the perceptual boundary means that despite sensory encoding noise, nearly all of the perceived locations are in the /ba/ regions of representational space (Fig. 1B). However, for repeated presentations of Stimulus 2, its location near a perceptual boundary means that sensory noise places some of the perceived locations in the /da/ regions of representation space (Fig. 1C).

The CIMS model assumes that auditory syllables and McGurk syllables are processed by the same perceptual mechanisms, resulting in predictable differences if Stimulus 1 and 2 are paired with an identical visual /ga/ in a McGurk A/ba/V/ga/ stimulus. For Stimulus 1, optimal cue combination produces an integrated representation that lies predominantly in the /ba/ regions of representational space, resulting in primarily “ba” percepts (Fig. 1D). For Stimulus 2, the integrated representation lies primarily in the /da/ region of representational space, corresponding to the McGurk fusion perception of “da” (Fig. 1E).

### Experiments 1 and 2: Finding strong and weak McGurk stimuli, with replication

To test these predictions, in Experiment 1 twenty productions of auditory /ba/ were recorded from a single female talker, in the expectation that natural variability in speech production would result in syllables of varying strength. Each different /ba/ was paired with a single visual /ga/ recorded from the same female talker and the resulting McGurk AbaVga stimuli were presented to participants. Across stimuli, there was substantial variability in the responses, ranging from 25% fusion responses to 65% fusion responses (Fig. 1F). Participants were also presented with unmanipulated control stimuli consisting of three different congruent audiovisual syllables (A/ba/V/ba/, A/da/V/da/, and A/ga/V/ga/). Participants responded accurately to the control stimuli, demonstrating that they were engaged in the task and not responding randomly (see Table 1 for data from control stimuli for all experiments).

**Table 1.** Each row provides the mean and standard error of the mean (SEM) accuracy across participants for the control stimuli (Congruent Performance) and the response percentages to the McGurk stimuli (first averaged across stimuli, then mean and SEM across participants).

| Exp | N | Congruent Performance |  |  | # Stim | Responses to AbaVga |  |  |
| --- | --- | --- | --- | --- | --- | --- | --- | --- |
| | | (mean ± sem) | | | | (mean ± sem) | | |
|  |  | ba | da | ga |  | ba | da | ga |
| 1 | 128 | 97 ± 1 | 87 ± 2 | 98 ± 1 | 20 | 26 ± 3 | 46 ± 3 | 28 ± 2 |
| 2 | 40 | 97 ± 1 | 94 ± 3 | 98 ± 1 | 2 | 37 ± 5 | 44 ± 5 | 19 ± 4 |
| 3 | 40 | 96 ± 2 | 80 ± 6 | 98 ± 2 | . | . | . | . |
| 4 | 40 | 99 ± 1 | 93 ± 3 | 97 ± 2 | 3 | 34 ± 5 | 46 ± 6 | 20 ± 4 |
| 5 | 44 | 98 ± 1 | 98 ± 1 | 88 ± 3 | . | . | . | . |
| 6 | 38 | 96 ± 3 | 86 ± 5 | 100 ± 0 | 2 | 41 ± 7 | 41 ± 5 | 18 ± 4 |
| 7 | 34 | 97 ± 2 | 77 ± 6 | 97 ± 1 | 4 | 35 ± 7 | 56 ± 7 | 6 ± 3 |

From the twenty different McGurk stimuli, we selected two stimuli at opposite ends of the distribution for further investigation, labeling them “S1” and “S2” by analogy with the modeling above. To ensure the stimulus differences between S1 and S2 were reliable, in Experiment 2 we attempted to replicate the results of Experiment 1, presenting only S1 and S2. Similar rates of McGurk fusion responses were observed (Fig. 1G; S1: 33% original *vs*. 28% replication; S2: 62% *vs*. 59%). A linear mixed-effects analysis on fusion responses showed a significant effect of stimulus [*t*(80) = -4.3, *p* = 10^−5^], but no effect of experiment [t(156) = -0.4, *p* = 0.66] or stimulus-by-experiment interaction [*t*(80) = -0.18, p = 0.86].

### Experiments 3 and 4: Comparing McGurk effect and clear auditory syllables

Next, we created a morphed stimulus that was a combination of S1 and S2. This morphed stimulus (labeled “S1.5”) allowed us to test two predictions of the CIMS model. First, CIMS predicts that if S1 and S2 lie at different locations in the one-dimensional representational space, then S1.5 should lie between them. Second, CIMS predicts that there should be a relationship between the perception of auditory /ba/ syllables presented alone and in combination with visual /ga/.

To test the first prediction, in Experiment 3 we presented the three auditory-only /ba/ components of S1, S1.5 and S2 to 40 participants. As expected, there was a decrease in the number of /ba/ percepts from S1 to S1.5 to S2 (Fig. 1H; 96% to 92% to 87%).

To test the second prediction, in Experiment 4, we presented the three stimuli (S1, S2, and S1.5, each consisting of a different auditory /ba/ paired with same visual /ga/) to 40 participants. As expected, there was an increase in the McGurk fusion percepts from S1 to the S1/S2 morph to S2 (Fig. 1I). Replicating the results of Experiments 1 and 2, S2 produced more fusion responses than S1 (54% *vs*. 38%; paired *t*-test, *t* = -2.4, *p* = 0.02). The S1.5 morphed stimulus produced an intermediate level of fusion responses (46%).

To determine if there was a stimulus-level relationship between syllable perception and the McGurk effect, we plotted the rates of McGurk perception for S1, S1.5 and S2 against the perceptual accuracy for the auditory component of each stimulus (Fig. 1J). To test for the linearity of this relationship, we compared two LMEs. One that assumed a linear relationship between stimuli (coded 0, 0.5 and 1) and one that allowed the stimuli to vary freely (categorical coding of stimuli). Comparing BIC between the two models yielded a better fit for the model assuming a linear relationship (BIC difference 9). For the winning model, the estimated slope across stimuli was 16 ± 6 [*t*(80) = 2.6, *p* = 0.01].

### Experiments 5 and 6: Comparing McGurk effect and noisy auditory syllables

The CIMS model predicts that adding noise to the sensory stimulus should broaden the distribution of perceived locations, which should differentially affect the different stimuli, with a bigger effect on the weaker S2 stimulus. To test this prediction, in Experiment 5 we added auditory noise to the auditory-only /ba/ component of S1 and S2 and presented them to 40 participants. This had the effect of decreasing the number of /ba/ reports as noise increased for both Stimulus 1 and Stimulus 2 (Fig. 1K). However, at all noise levels, there were fewer /ba/ reports for the weaker Stimulus 2 than the stronger Stimulus 1, as predicted by CIMS. An LME with fixed factors of noise (entered as SNR, with clear speech set to +12dB), stimulus, and their interaction along with random effect of subject yielded significant main effects for stimulus [*t*(484) = 7.0, p = 10^−11^], noise [*t*(484) =18.9, *p* < 10^−16^] and the interaction [*t*(484) = -4.8, *p* = 10^−5^]. Compared with clear speech, at -12 dB, the number of /ba/ reports decreased only slightly for S1 [99% *vs.* 98%; paired *t*(43) = -0.8, *p* = 0.42] but decreased greatly for S2 [98% *vs.* 53%; *t*(43) = -8.0, *p* = 10^−10^].

### Section 2: Participant Variability

There are two sources of participant variability in CIMS. The first source is individual differences in sensory encoding noise. Observer 1 might precisely encode speech, creating a narrow distribution of perceived locations in representational space, while Observer 2 might imprecisely encode speech, creating a broad distribution (Fig. 2A). When presented with an auditory /ba/ near the perceptual boundary, the precise representation of Observer 1 will result in mainly /ba/ percepts while the imprecise representation of Observer 2 will blur many of the estimates into the /da/ regions of representational space (Fig. 2B).

**Figure 2.**
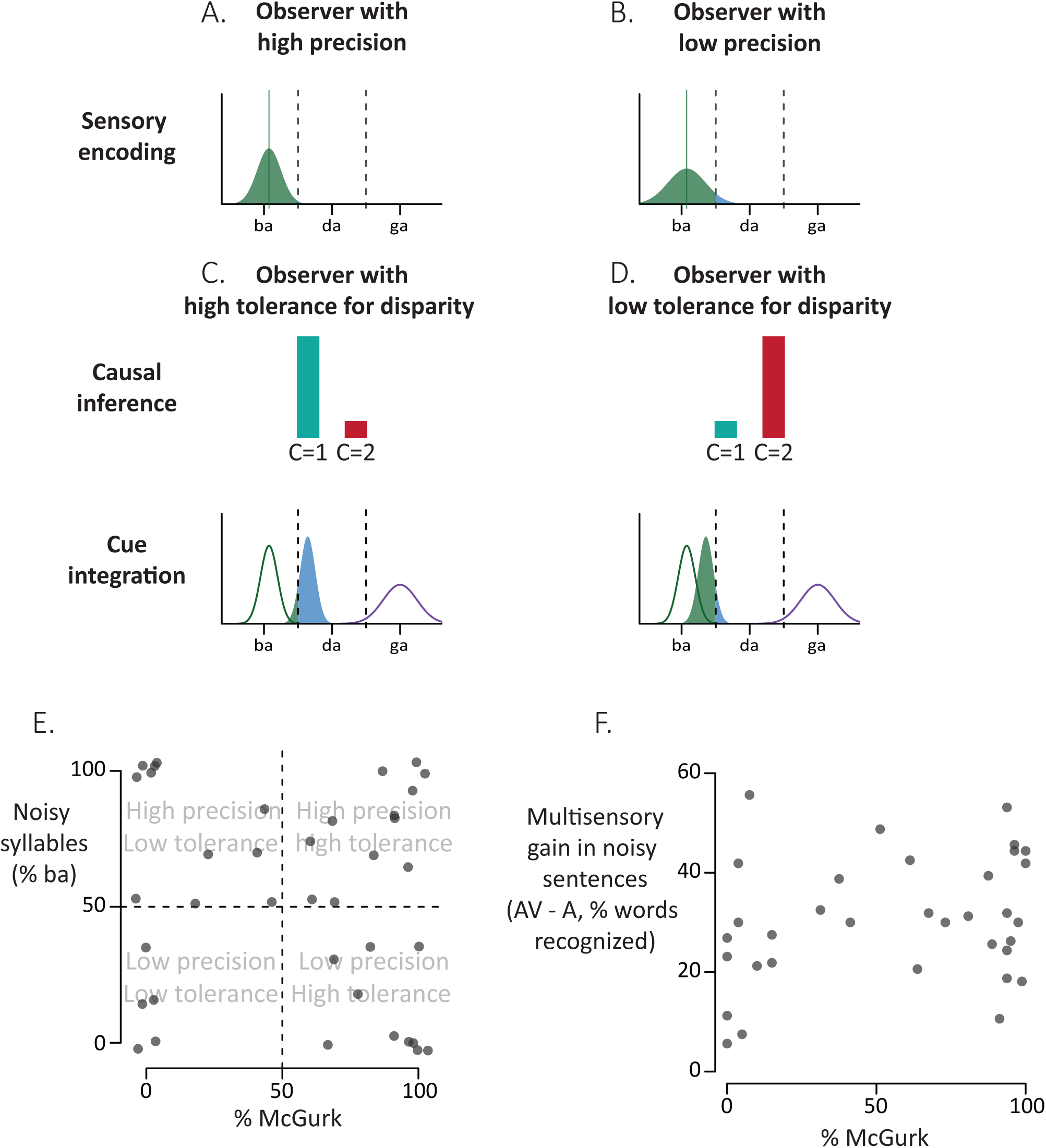
The CIMS model applied to participant variability. (A) Variability in speech perception between participants can arise from individual differences in sensory encoding. For an observer with high precision, there will be a narrower distribution of perceived locations for a given stimulus (shown for Stimulus 1 from Fig. 1A). (B) For an observer with low precision, there will be a broader distribution of perceived locations for a given stimulus (shown for Stimulus 1 from Fig. 1A). (C) Even if encoding precision is identical, variability in speech perception between participants can also arise from individual differences in causal inference. For an observer with high tolerance for disparity presented with a McGurk stimulus consisting of auditory /ba/ and visual /da/, the observer infers that the auditory and visual cues arise from a single talker (C = 1, illustrated as relative heights of C = 1 and C = 2 bars). Optimal cue integration reflects this inference so that the integrated representation lies between the auditory and visual representations, with most percepts falling in the /da/ region of representational space (blue shaded region, high rates of McGurk,). (D) An observer with low tolerance for disparity infers that the auditory and visual speech in the McGurk stimulus arises from two different talkers (C = 2). Optimal cue integration reflects this inference so that the integrated representation is weighted by the auditory-only representation, with most percepts falling in the /ba/ region of representational space (green shaded region, low rates of McGurk). (E) Rates of McGurk and accuracy of noisy auditory syllable presentation across participants, one symbol per participant, *r*(36) = -0.09, *p* = .60. The noisy syllable measure is the accuracy of perception of stimulus S2 with -12 dB noise added from experiment 5 and the rate of McGurk perception is from experiment 6. Participants were distributed across quadrants with all combinations of low and high sensory encoding precision and low and high tolerance for audiovisual disparity. (F) Rates of McGurk and multisensory gain during noisy sentence perception across participants in experiment 7, one symbol per participant; *r*(31) = 0.261, *p* = 0.14. The audiovisual gain measure is the percentage of keywords understood during perception of noisy sentences with the talker visible minus the percentage of keywords understood during perception of auditory-only noisy sentences.

The second source of participant variability in CIMS is individual differences in causal inference. Multisensory integration is only beneficial if the cues to be integrated were caused by the same real-world event (a single cause, C = 1); integrating cues generated by two different real-world events (C = 2) worsens perception. For presentation of clear or noisy syllables, all cues indicate that C = 1, reducing any participant variability in causal inference. In contrast, McGurk stimuli are created by dubbing incongruent auditory and visual syllables, creating a conflict between the temporal synchrony and spatial coincidence of the auditory and visual syllables (which suggest that C = 1) and the content disparity between the heard speech and the viewed mouth movement (which suggests that C = 2). For observers with a high tolerance for content disparity (leading them to infer that C = 1) optimal cue combination will more strongly weight the integrated representation of auditory and visual speech, usually resulting in the illusory McGurk percept. For observers with a low tolerance for content disparity (leading them to infer that C = 2) optimal cue combination will more strongly weight the reliable auditory-speech representation, usually resulting in a percept corresponding to the auditory token.

### Experiments 5 and 6: Participant variability in McGurk effect and noisy syllable perception

To test these ideas, we examined the performance for those participants who participated in both experiment 5 (where they were presented with noisy syllables) and experiment 6 (where they were presented with McGurk stimuli). To prevent floor or ceiling effects, the measure for noisy syllable perception was the accuracy of perception of stimulus S2 presented at the -12 dB noise level in experiment 5. There was substantial variability in perception of the McGurk effect, with rates ranging from 0% to 100% and in perception of noisy syllables, with accuracy ranging from 0% to 100%. However, across participants there was low correlation between the two values [Fig. 2E; *r*(36) = -0.09, *p* = .60]. Participants were found in all quadrants of the plot, modeled by CIMS as participants with low or high encoding noise and low or high tolerance for audiovisual disparity in the causal inference judgment.

Next, we considered all noise levels across participants by comparing two LMEs: the original LME fit to the noisy syllable data (dependent variable accuracy; fixed effects of stimulus, noise level and their interaction; random effect of subject) and a second LME that additionally contained subject-level McGurk perception (per stimulus). Comparing BIC between models, we found that knowing McGurk responses did not explain additional variance in noisy-syllable perception (BIC difference 11). Importantly, this absence of participant-level relationships between McGurk and noisy syllable perception occurred despite the presence of a stimulus-level relationship: S1 was more accurate than S2 (77% *vs.* 48%, across all noise levels), but S2 produced more McGurk responses (61% *vs.* 39%).

### Experiment 7: Participant variability in McGurk effect and noisy sentence perception

To assess whether there was an across-participant correlation for more complex forms of speech, in Experiment 7 we compared perception of the McGurk effect and perception of noisy sentences in 33 participants. There was substantial variability in the rate of perceiving the McGurk effect across different participants, from 5% to 95% (Fig. 5A). For noisy sentences, participants correctly reported 14 out of 160 (9%) of keywords for auditory-only sentences but correctly reported 62 out of 160 (39%) of key words for audiovisual sentences, a 30% improvement [AV - A, (paired *t*(33) = 14.0, *p* = 10^−15^)]. However, there was substantial variability in the visual benefit across participants. The participant with the lowest benefit recognized only 6% more words when viewing the face of the talker while the participants with the highest benefit recognized 50% more words. For consistency with previous reports, in addition to the multisensory gain (AV - A) we also calculated a visual enhancement index for each participant, VE = [(AV - A)/(1 - A)] (Van Engen et al., 2017). The mean visual enhancement was 34% (range 6% to 63%).

Next, we compared the two axes of individual variability. There was low correlation between rates of McGurk effect and multisensory gain, *r*(31) = 0.261, *p* = 0.14 (Fig. 2F); between rates of McGurk effect and the visual enhancement index, *r*(31) = 0.256, *p* = 0.14; and between rates of McGurk effect and auditory-only performance, *r*(31) = -0.06, *p* = 0.73. As in the syllable data, participants fell in all 4 quadrants of the plot, with high and low McGurk susceptibility and high and low multisensory gain.

## Discussion

In the present study, we used natural variation in the production of the syllable /ba/ to understand the relationship between the McGurk effect and other speech perception tasks. Pairing twenty unique auditory /ba/ syllables with a single visual /ga/ produced McGurk stimuli that elicited reliably different levels of McGurk perception across large groups of subjects. For two stimuli that differed substantially in their McGurk strength, the auditory /ba/ that was less effective at evoking the McGurk effect was recognized more accurately in both clear and noisy auditory-only perception tasks. The stimulus-level relationship between McGurk and auditory-only perception was well-described by the CIMS model.

Across participants, the CIMS model also provided a straightforward explanation of why perception in two seemingly related behaviors, McGurk perception and perception of noisy speech, are not highly correlated. In audiovisual speech in noise tasks, individual differences primarily arise from differences in sensory encoding (how well one understands the individual speech tokens). In contrast, the disparity inherent in the McGurk stimulus means that causal inference judgments can create individual differences in perception even in individuals with identical sensory encoding.

### Stimulus differences in the McGurk Effect

Our study was motivated by recent findings that have raised questions about the utility of the McGurk effect as a tool for understanding everyday speech perception. Basu Mallick and colleagues (Basu Mallick et al., 2015) reported high variation in the McGurk effect across different stimuli (rates ranging from 17% to 58%) and participants (rates from 0% to 100%). While a careful study of many acoustic and visual properties of McGurk stimuli showed that taken together they accounted for about half of the variability in the frequency of the effect across stimuli and participants (Jiang and Bernstein, 2011), it is not clear if the factors contributing to variability in the McGurk effect are relevant for everyday speech perception.

The stimuli in the Basu Mallick study were culled from different sources and contained different talkers, different views of the face and upper body, different native languages, and different video and audio quality. To better control these factors, in the present study we created a new corpus of McGurk stimuli that were all recorded from the same talker within a short time span and had the same visual component, making them closely matched for auditory and visual properties. Nevertheless, there was still substantial variability, and this variability was related to the perception of the auditory component of each stimulus in the direction predicted by the CIMS model. Auditory /ba/ tokens modeled as being more prototypical were more likely to evoke a /ba/ percept when presented alone, either with or without added noise, and were less likely to evoke a McGurk fusion percept when paired with a visual /ga/.

Speech production is known to be variable in both articulation and acoustics (Holmberg et al., 1994; Whalen et al., 2018). Although we did not quantify physical differences between stimuli, such as voice onset time (Wood, 1976) our findings show that variability in production leads to variability in perception that is similar for McGurk and everyday speech. If both types of speech are processed using similar computations (as shown by the stimulus-level relationship between them) then the McGurk effect can be a useful tool for interrogating everyday speech perception.

### Relating the McGurk Effect to other speech perception tasks

A second critique of the McGurk effect is the concern that variability in McGurk perception is unrelated to variability in perception of everyday speech (Alsius et al., 2018). Van Engen and colleagues (Van Engen et al., 2017) found low (and non-significant) correlations between McGurk effect and visual enhancement of noisy speech, while Brown and colleagues (Brown et al., 2018) used a larger sample size to find a low (but significant) correlation between McGurk effect and lipreading accuracy. Other studies have also found complex relationships between different measures of auditory, visual and audiovisual speech perception (Grant and Seitz, 1998, 2000; Sommers et al., 2005; Strand et al., 2018).

Like many fields of inquiry, research in speech perception has been muddled by the fallacy that statistical significance signals truth (Ioannidis, 2005). Stimulus and participant variability are high and sample sizes are usually small (Magnotti and Beauchamp, 2018). This can result in seemingly contradictory results, in which one study finds that two processes are significantly correlated while another finds that they are not. However, the correlation values themselves may not be significantly different between the two studies (Gelman and Stern, 2006).

A lack of correlation between the McGurk effect and other measures of speech perception is surprising only if audiovisual speech integration is assumed to be a single, unitary phenomenon. The CIMS model parcellates audiovisual speech integration into discrete stages, each of which serves as a source of individual variability. Perception of noisy speech and lipreading do not require causal inference, as all of the cues indicate that the speech arises from a single talker. In contrast, perception of the McGurk effect requires a causal inference judgment because of the conflicting cues from temporal synchrony (which suggests that C = 1) and content incongruity (which suggest that C = 2). In the CIMS model, individual differences in noisy syllable perception arise from differences in sensory encoding, while individual differences in the McGurk effect include the additional factor of differences in tolerance for audiovisual disparity.

Our finding of observers with all combinations of low and high sensory encoding precision and low and high disparity tolerance shows that these individual differences are not correlated within individuals. While the CIMS model is agnostic with respect to neuroanatomy, BOLD fMRI and modeling suggest that there are anatomical dissociations between brain areas responsible for sensory encoding and those responsible for causal inference judgments (Rohe and Noppeney, 2015; Cuppini et al., 2017) making it reasonable that individual differences in one computation are uncoupled from individual differences in the other. Findings that the McGurk effect shows different neural signatures than congruent audiovisual syllables (Erickson et al., 2014; Moris Fernandez et al., 2017; Sánchez-García et al., 2018) has been used as evidence that the McGurk effect is processed differently than everyday speech. The CIMS model clarifies that McGurk stimuli place higher demands on neural circuits underlying causal inference judgments than does processing congruent audiovisual syllables. Therefore, the CIMS model predicts that activity in causal inference circuits should be observed for both the McGurk effect and some everyday conditions such as multiple audiovisual talkers but not for congruent speech from a single talker.

Although the McGurk effect is an artificial lab creation, causal inference is an important step in perception whenever observers are confronted with multiple possible sources for sensory cues, such as multiple talkers speaking at once (Kording et al., 2007; French and DeAngelis, 2020). For instance, in the ventriloquist illusion, spatial disparity cues suggest that C = 2, since the voice of the puppeteer and the dummy arise from different spatial locations. However, synchrony cues suggest that C = 1, since the dummy’s mouth moves in time with the auditory speech (reviewed in Bruns, 2019). Evidence suggests that cross-subject variability in the tendency to bind audiovisual signals is found across a range of tasks, and that these differences are stable across time but are task-specific (Odegaard and Shams, 2016).

We replicated the findings of Van Engen and colleagues (Van Engen et al., 2017) finding low correlation between the McGurk effect and noisy sentence perception tested with a variety of measures. As pointed out by Van Engen and colleagues, perception of noisy sentences calls on many cognitive processes in addition those required for syllable or McGurk perception (Davis and Johnsrude, 2007). Contextual information is thought to be modulated by top-down projections from frontal cortex, a different set of brain areas from the networks responsible for sensory encoding and causal inference judgments (Peelle and Sommers, 2015; Gau and Noppeney, 2016; Tuennerhoff and Noppeney, 2016; Cope et al., 2017). Therefore, the top-down processing required for sentences provides yet another opportunity for cross-participant variability to arise, and it is entirely possible for this variability to be uncorrelated with individual differences in either sensory encoding or causal inference.

### Conclusions

The CIMS model provides a straightforward explanation for our findings of a high stimulus-level correlation between McGurk and everyday speech perception (explained by a common representational space for both kinds of speech) but a low participant-level correlation between the two types of speech (explained by separate axes of individual differences in sensory encoding and causal inference judgments). “Audiovisual speech perception” (or “multisensory integration”) is unlikely to be a unitary phenomenon easily captured by a single behavioral measure (Soto-Faraco and Alsius, 2009). Many measures of everyday speech perception show high individual variability and low correlation across participants. Therefore, suggestions to retire the McGurk effect because it shares these attributes may be premature (Alsius et al., 2018; Rosenblum, 2019). Instead, we suggest the alternative approach of creating an explicit model of the speech perception process of interest and using the model to guide selection of the appropriate behavioral measures.

## Acknowledgments

This research was supported by NIH R01NS065395.

## Notes

### Competing Interest Statement

The authors have declared no competing interest.

